# Automated, Stress-Free, and Precise Measurement of Songbird Weight in Neuroscience Experiments

**DOI:** 10.1101/2024.10.05.616794

**Authors:** Yuval Bonneh, Avishag Tuval, Ido Ben-Shitrit, Lilia Goffer, Yarden Cohen

## Abstract

Monitoring the health and well-being of research animals is essential for both ethical and scientific purposes. In songbirds, body weight is one of the main indicators for their overall condition, yet traditional weighing methods can be intrusive and stress-inducing, which could decrease their song rate. We developed a novel, automated system designed to continuously monitor the weight of untethered and tethered birds without disrupting their natural behavior in neuroscience experiments. We used the system to track weight fluctuations in six canaries over several weeks, revealing physiological patterns such as overnight weight loss, with one bird losing approximately 5.17% of its body weight during a 9.5-hour period of inactivity. Our system’s high sensitivity detected weight changes below 1% of body mass, validating its reliability for long-term studies. Control experiments confirmed that weight fluctuations observed were physiological rather than due to equipment deviations. By eliminating the need for manual handling, this system offers a non-invasive, hands-free approach that reduces stress and improves the accuracy of health assessments. This study demonstrates the system’s potential for expanding research on how environmental factors, diet, and other variables influence bird physiology and behavior. Future applications could integrate additional health metrics, providing a more comprehensive understanding of animal welfare in neuroscience and behavioral studies.

## 1 Introduction

Songbirds, including canaries and zebra finches, are excellent animal models for studying the neural basis of vocal learning and motor sequence generation [3, 6, 9]. These studies often necessitate surgical procedures, such as electrode or probe implantation in the brain, followed by longitudinal observations that may extend over days to months, conducted on tethered birds in acoustic chambers [16]. Such studies require stringent protocols for housing, husbandry, and monitoring to ensure the birds’ well-being while maintaining conditions conducive to singing behavior in the lab [12]. Healthy birds are more likely to produce high quality data, and therefore this kind of protocols is not only crucial for the birds, but also beneficial for the researchers, seeking reliable data that represents natural behaviours.

Body weight is a critical indicator of health and well-being in laboratory animals, offering an objective measure of physiological status [10]. Weight loss can reflect decreased appetite as a consequence of distress, fear and pain, but it can also indicate progression of a chronic disease reflecting deterioration with an increasing burden for the animals [13]. Significant deviations from baseline weight, particularly a loss of 20% or more, are used as humane endpoints across species [10]. For birds, monitoring body weight can provide valuable insights into their overall condition, especially when tracked over time. Research has shown that adult birds tend to lose weight following changes in husbandry [16], and experience weight fluctuations during reproductive and molting cycles [2]. Birds can even show daily and seasonal variations: Daily weight fluctuations of 5 to 10% are common, typically being highest in the late afternoon and lowest after fasting at night [8]. Seasonal variations also play a role, with birds often being heavier in the winter to survive harsher conditions [2]. Some birds exhibit changes corresponding to seasonal rhythms even when kept under constant laboratory conditions [16], and hormonal regulation of these processes has been demonstrated [7]. Thus, routine weight monitoring during neurophysiological studies in songbirds could yield insights into hormonal states among other factors influencing their singing behavior.

Despite the potential benefits of weight monitoring, it is not commonly reported in birdsong research. Manual weighing methods, such as placing birds in cloth bags for measurement [5], can induce stress and affect song rate in laboratory conditions. Therefore, in experiments where birds are housed in acoustic chambers for song recording, handling of birds is minimized to avoid altering song output. As an alternative to weighing, Yamahachi et al. showed that song rate can indicate stress levels and demonstrated that singing more than several hundred motifs per day suggests effective stress coping in zebra finch [15]. This approach proved to be very reliable in a range of stressors, for example post-surgeries, after tethering birds to experimental setup, and in longitudinal song recordings in isolation, but it is specific to male zebra finches and relies on their stereotyped behavior -their stereotyped song motifs and their stable daily song rate. In many other situations, song rate remains an unreliable measure of well-being. A stereotyped song motif and a stable singing rate cannot be used in other songbird species, for example in various sparrow species and in canaries that have seasonal changes in singing rate and highly complex songs with no countable motifs. Song cannot be used as a reliable readout also in zebra finches: in females, who do not sing, or after surgical procedures targeting brain areas in the *song premotor system* that could decrease the bird’s singing behaviour regardless of the animal’s well being [12]. Furthermore, a song-rate measure requires a full day of recording before motifs can be counted. In contrast, weight measuring gives an instantaneous readout and can be sensitive enough to show diurnal effects. In sum, the rate of song motifs is a reliable readout of well being in a limited set of experiments on male zebra finches. In many other cases, additional well-being metrics are essential.

To address these challenges, we developed an automated, stress-free system for monitoring the weight of freely behaving songbirds, both tethered and untethered, in order to detect weight loss as a marker for their well-being. As automated systems have been previously developed for mice [11], Our system is tailored to avian anatomy and compatible with birds housed in acoustic chambers. This system enables monitoring the weight of up to 8 birds simultaneously, continuously (24/7), and without interaction, minimizing the effects on their natural behaviour such as song rate. We demonstrate that our system allows reliable tracking of individual birds’ weight over weeks and that it is sensitive enough to observe the gradual decrease in body weight overnight -validating its potential to monitor the well being of songbirds. To help test this possibility and its benefits beyond the specific use in our lab, we also provide the complete mechanical, electronic, and software designs of our system.

## 2 Methods

### 2.1 System design

#### Overview

The Scale System is a non-invasive setup designed to monitor the weight of multiple, freely-behaving birds directly while in their cage, in order to detect weight loss as a marker for their health condition. The weighing is conducted by placing a load-cell-based weighing device (scale) in the bird’s cage. This device is connected to a load cell amplifier (NAU7802, Sparkfun) which helps convert and amplify the signal from the load cell to an I2C signal, readable with an Arduino micro-controller from which we can extract the weight in grams. The data collection is operated using a python script that runs continuously on a Raspberry Pi minicomputer. This script receives data from the Arduino, and stores it as ‘.csv’ files. Each *Scale System* is capable of connecting up to 8 weighing devices per micro-controller with the help of another breakout board (MUX), which enables communication with multiple I2C devices simultaneously.

#### Scale design and technical information

The Scale device (Fig. 1A) is comprised of a 3D printed Perch (150 mm long, 12 mm dia. made out of Delrin, scratched for better grip), a 400 g custom-shaped steel cylinder as a counterweight, and a 500 g load cell (SEN14728, Sparkfun). The load cell is connected to the perch on one side and to the steel weight on the other with M3 screws (two on each side) so that the perch floats 8 mm above the surface, avoiding complications with tethered birds’ cables. The steel weight is carved from a 50 mm dia., 16 mm tall cylinder to form designated gaps for the load cell to latch to and for the load cell wires to pass under the weight (Fig. 1B). For our convenience, the ‘base’ of the weight is designed to be 16 mm tall and weigh ∼ 200 g, with two additional 100 g plates of the same diameter with a hole in the center for easily screwing them one on top of the other, potentially getting 300-or 400-gram weight to counter the weight and momentum of a bird flying on to the perch, keeping the scale stable (full design is avialable at https://github.com/NeuralSyntaxLab/Bird-Scale-Methods-Article/tree/main/Scale%20design). Canaries are considered to weigh somewhere between 15 and 25 grams [4], and the perch weighs 25 g, so the total possible weight upon the load cell is approximately 50 g. Regardless, we chose to use a load cell with a capacity of 500 g and precision of 0.1 g to withstand the force of a bird flying onto the perch with speed while maintaining accurate measures.

**Figure 1.**
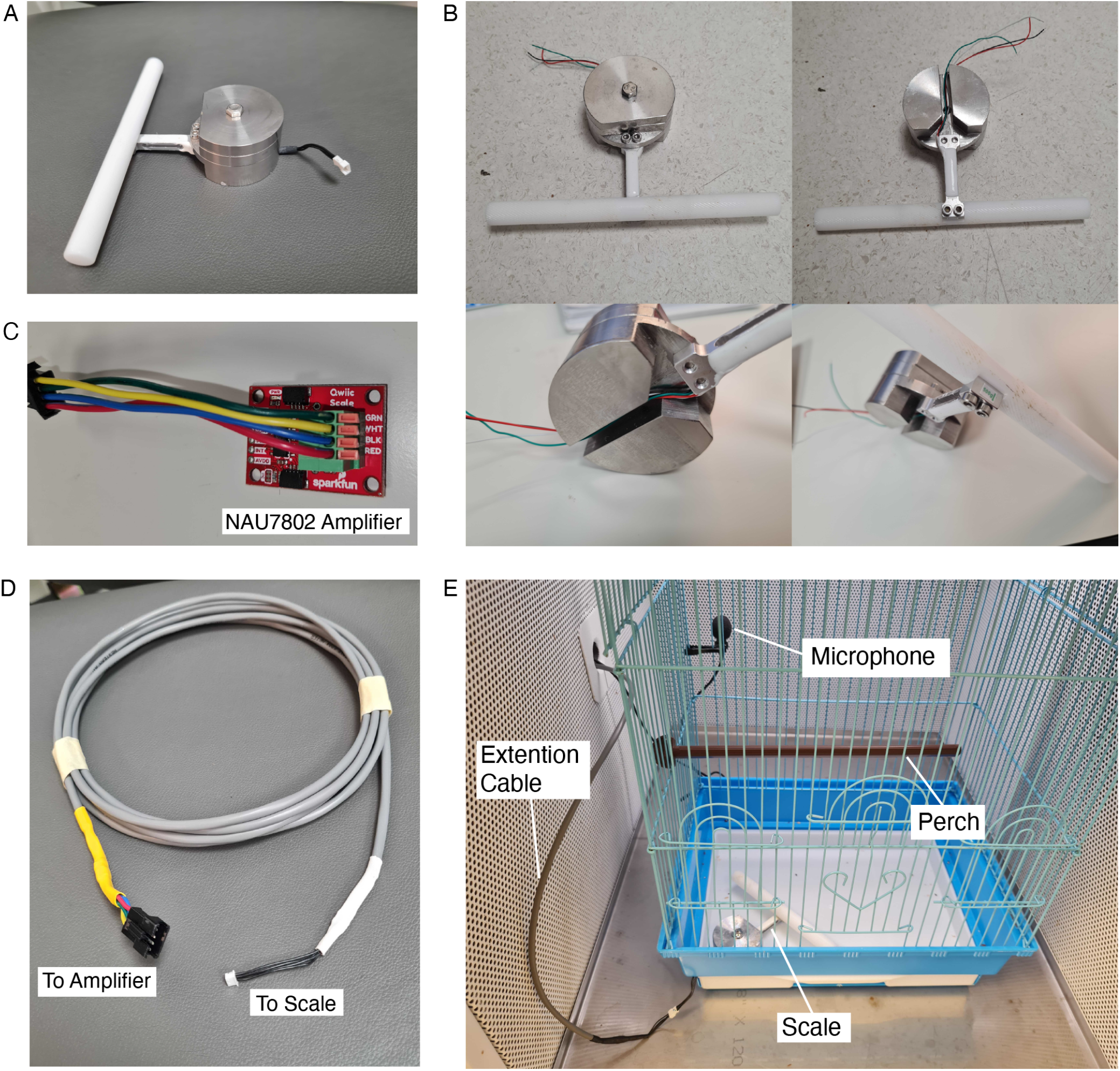
Scale system components and setup design. **(A)** The scale device. **(B)** Scale assembly nuances depicted, including an overview, underview, and angles showing the connections between the load cell, the perch and the steel weight. Notice the gap under the steel weight where the load cell cables go. **(C)** Image of the load cell amplifier (nau7802 board). Notice the labeled spring terminals where the green, white, black and red wires of the load cell connect. In this image, green, yellow, blue and red wires of a JST-SM 4-PIN Pigtail connector’s male plug are inserted to the sockets. **(D)** Image of the Communication Cable, soldered to the Circuit Picoblade plug 425mm female connector on the scale side, and to a JST-SM 4-PIN Pigtail connector male plug that will match the one that is shown in figure 1c inserted to the amplifier. **(E)** Image of the scale setup inside an Acoustic chamber, fixed to the bottom corner of the birdcage. Notice the 4-pin connector sticking out of the scale in the corner of the cage, connected to the communication cable’s, which spreads outside of the acoustic chamber through the designated hole in the side.

#### Technical design

The load cell’s 4 wires (positive and negative signals – green and white, power – red, ground – black) are soldered to either plug (male or female) of a 4-pin connector (Circuit Picoblade Male-to-Female plug 425 mm) as the other plug is soldered to one end of a 4-core *Extension Cable* (Fig. 1D). This way, easy insertion and extraction of the scale in and out of the birdcage is enabled, before connecting it to the load-cell amplifier using the long extension cable, so that eventually the load cell’s 4 wires are connected to the corresponding labeled spring terminals on the amplifier (Fig. 1C). This amplifier is an Analog-to-Digital converter (ADC) with built in gain and I2C output. This device amplifies the signal from the load cell and converts it to I2C signal that an Arduino micro-controller can read. We can also use this signaling method to read data from multiple scale devices simultaneously, with the help of another breakout board (Qwiic MUX, Sparkfun), which enables communication with up to 8 I2C addresses. Therefore, the extension cables are used to allow connecting multiple scales scattered around the room to one control unit (MUX, Arduino and Raspberry Pi). The load cell amplifiers are each connected to the MUX breakout board, which is connected to an Arduino UNO micro-controller using designated cables (Flexible Qwiic cables, Sparkfun). The red-, black-, blue- and yellow-colored wires of this cables are connected to the 3.3V (power), Ground, SDA (serial data) and SCL (serial clock) pins in the Arduino UNO accordingly (The SDA and SCL pins are used for I2C communication). The Arduino UNO is loaded with a specialized Arduino code using Arduino IDE. This code is designed for the Arduino to run at a baud rate of 9600 for communicating with the Raspberry Pi and transmitting weighing data from up to 8 scales within one second. The sampling rate of the load cell amplifier is set to its maximum sample rate of 320 samples per second and each scale reading is considering an average of 8 samples when calculating the weight in grams. These sampling rates produce an average operation time of ∼ 200 ms for acquiring and sending precise weight data from 8 devices sequentially.

#### Calibrating the scales

The load cell’s raw data measures a change in electric signal, so converting this raw data into accurate weight in grams, requires all scale devices to be calibrated before starting to record the data. The calibration process takes place on the Arduino IDE platform, using a specialized Arduino code which we developed, that uses built in functions from the Sparkfun QwiicScale library. In the process the system calculates the *Zero Offset* to be the number that resets the raw data to 0 when there is nothing on the load cell (except for the perch). Then, an item of known weight is loaded on the perch and its weight in grams is pushed to the system, that calculates the *Calibration Factor* as the division of the raw value and the actual weight of the item. These two calibration values are stored in dedicated locations within the Arduino’s Non-Volatile Memory using the EEPROM library, so that in fact, there are designated memory location indices that stores the *Zero Offset* and *Calibration Factor* for every MUX channel.

Upon connecting a new scale device to a MUX channel, the user needs to re-calibrate it to store the new calibration values in place of the old ones. Whenever new raw data from a MUX channel is measured, values from its corresponding storage location are pulled in order to convert the raw value to an actual weight value according to Eq. 1

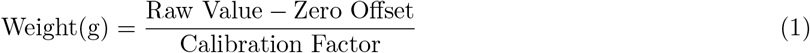

### 2.2 Experimental Procedures

#### Ethics declaration

All procedures were approved by the Institutional Animal Care and Use Committees of the Weizmann Institute of Science (protocol numbers: 08891223-1, 02110223-1, and 01850223-1).

#### Birds

A total of 6 birds were monitored for their weight using the scale system. Five of them were housed in 32*25*40 cm birdcages, placed inside acoustic chambers for 18 days while their singing was recorded for other experiments. These birds had two other perches in their cage for access to food and water as a part of the general lab procedures for housing birds. The other bird was housed in a specialized cage with dimensions of 32*25*32 cm and was being monitored while participating in another experiment of calcium imaging, thus a 2.5 g miniscope was mounted on its head. All birds were provided with a sufficient supply of food and water. Once a day the doors of all acoustic chambers were opened for the birds to socialize with each other for a duration of 1-2 hours.

#### Data collection

Weight data from all birds was continuously recorded at a frequency of 1 Hz by a python script running on the Raspberry Pi, which receives a list of 8 data points, each referring to one weight reading from a different scale, every second. This data is organized to match each of the weights to the relevant bird, and then saved in .csv format with the accumulated times (*‘date hh:mm:ss’* format) and weights (grams) for each bird.

#### Data analysis

Weight data from 4 birds was analyzed, as two were excluded for not standing on the scale. Initially, the data was cleaned from baseline, ‘off-scale’ values (*<* 10 g) and outliers (*>* 30 g). Then, Two methods were used for the extraction of reliable data points in order to estimate the measured weight: The first method considered sequences of non-baseline data that are longer than 10 seconds, as a reliable sequence. Then, the mean value of each sequence was calculated. The second method considered reliable data points as the mean values (calculated using a rolling windows with a window size of 100 samples) of rolling windows with lower standard deviation (*<* 1 g), and the mean weight per day was calculated based on these reliable data points. Further analysis using the first method found a few, very long sequences, where the bird stayed on the scale all night. For these sequences a linear regression was calculated to assess the bird’s weight loss over the night. All data analysis was done using in-house developed python code.

## 3 Parts List

A list of all the parts used and where to get them is downloadable through this link: https://github.com/NeuralSyntaxLab/Bird-Scale-Methods-Article/blob/main/Scale_system_parts_list.xlsx

## 4 Results

A scale system was inserted to the cages of 6 canaries (c.f. Fig. 1E), five of which were free to move around in their cage as their singing was recorded for other experiments. The other canary was housed in a different cage, free to move but tethered with a specialized flex cable to a rotating motor on the ceiling of the cage, as a 2.5 g miniscope was attached to its head for calcium imaging data acquisition. Weight data from the first 5 canaries was collected continuously for 18 days, and the tethered canary’s weight was monitored on-and-off for a few months, as the main focus of that experiment was the calcium imaging. Daily supply of food and water was given to all canaries, and once a day for 1-2 hours the acoustic chamber doors were opened for socializing. Another perch was available for the birds to stand on for consuming food and water from the designated apparatus for all canaries except for the tethered one. This means that these birds had perching options other than the weight measuring device -a point that we revisit in the Discussion section.

The weight data from one canary measured over 18 days shows numerous occasions per day where the canary stood on the scale for long enough in order to achieve reliable data points (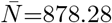*min*. = 63). After cleaning the data from baseline and outliers (see Methods), we calculated the mean weight using a rolling window (considered as one data point), with a window size of 100. The standard deviation of each window was also calculated, and only data points calculated from windows with a standard deviation lower than 1 g, also referred to as reliable data points, were included. Fig. 2 shows the measured weight over time for that canary during the entire 18-days period (reliable data points highlighted in orange), as well as a bar plot with the mean weight every day, calculated by averaging the values of all reliable data points within the same day. These findings indicate that the bird indeed stands on the perch frequently. The stability of weight readout suggests that the bird was maintaining a plane of controlled weight which indicates its physiological well being. To validate this possibility, we carried a separate control experiment, further discussed later. Its results are shown in Fig. 4C. This control measurement show that the scale is reliably maintaining its readout with fluctuations of 0.05 g -much smaller than the weight changes in Fig. 2.

**Figure 2.**
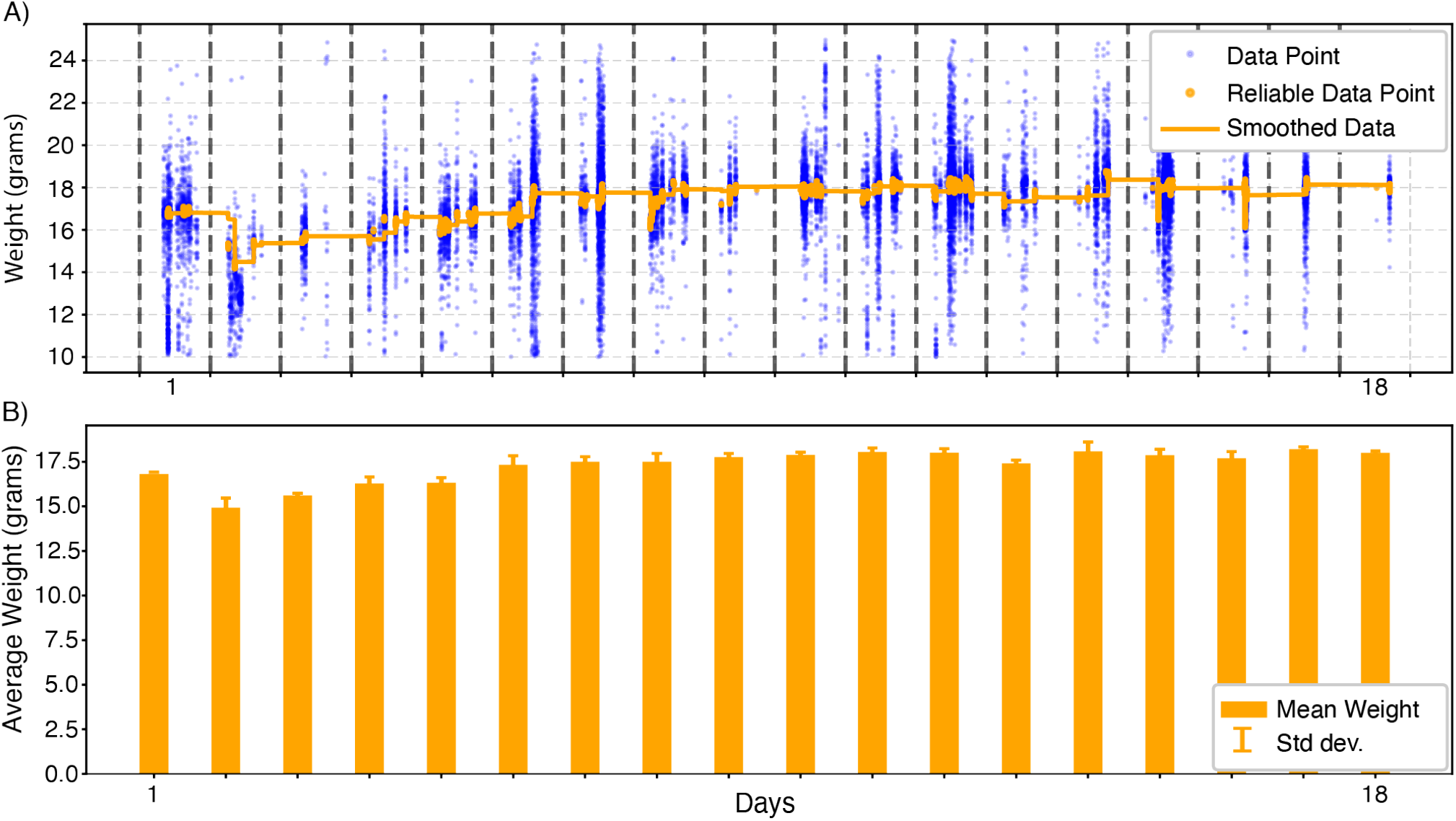
Average weight per day over 18 days of continuous measurements. **(A)** Data points of measured weight (grams) over time (blue dots) and calculated reliable data points (orange dots), as the orange line shows the reliable data smoothed. Vertical dashed lines mark day shifts. **(B)** The orange bars show the average weight per day, calculated from reliable data points. Error bars on top of each bin shows standard deviation among all reliable data points within that day.

**Figure 3.**
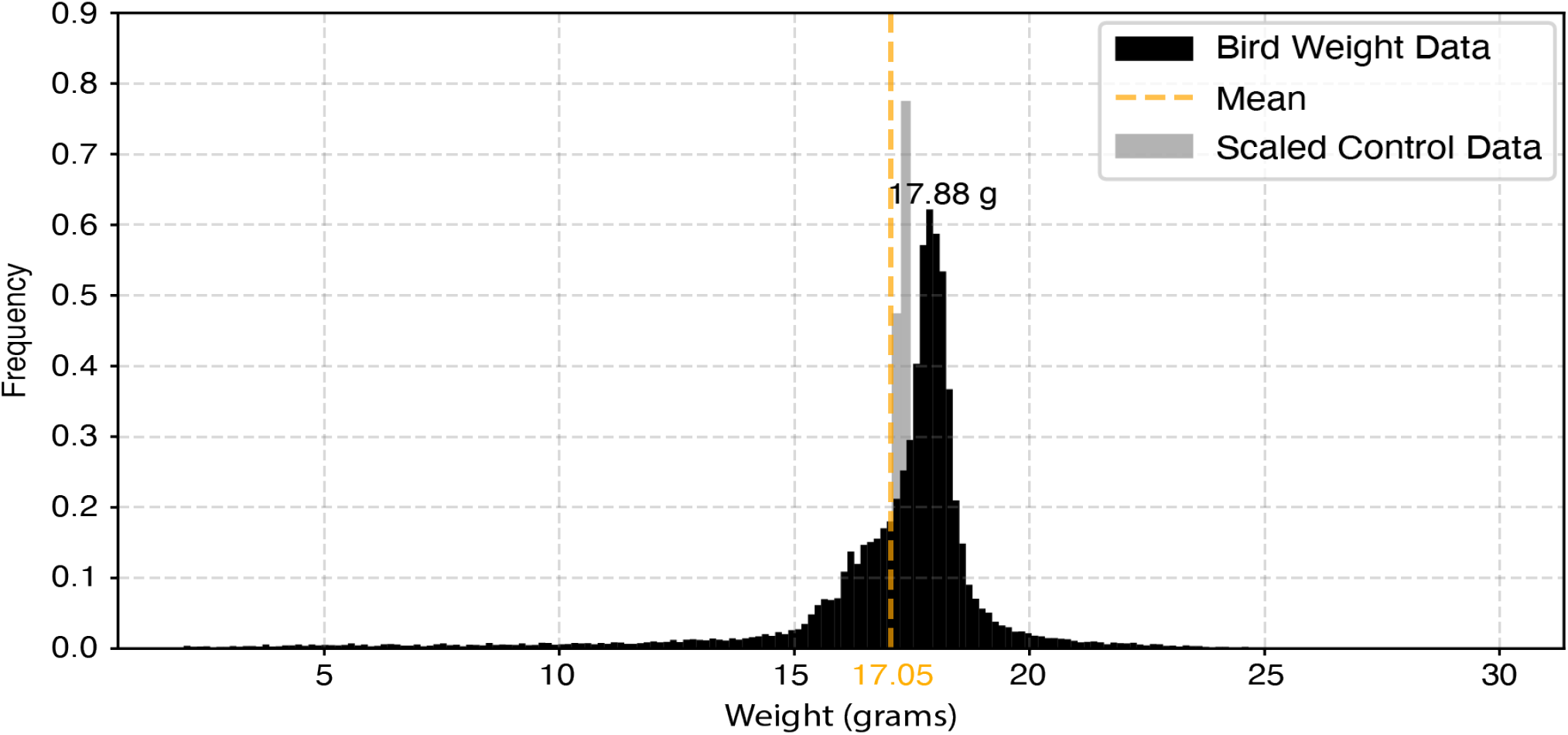
Density distribution histogram of all measured weight data points throughout 18 days of monitoring, cleaned from baseline ‘off-scale’ values lower than 2 g (black, *n* = 44896), binned to 0.1 g bins. This distribution shows the most frequent value of 17.88 g. The orange vertical dashed line shows the mean value of this dataset (17.05 g). Data from the control experiment (*n* = 71916) was binned with a bin size of 0.2 g, shown here in grey. The bins are scaled by 0.25 from their actual density proportion in order to normalize them to the dimensions of the canary data density distribution.

**Figure 4.**
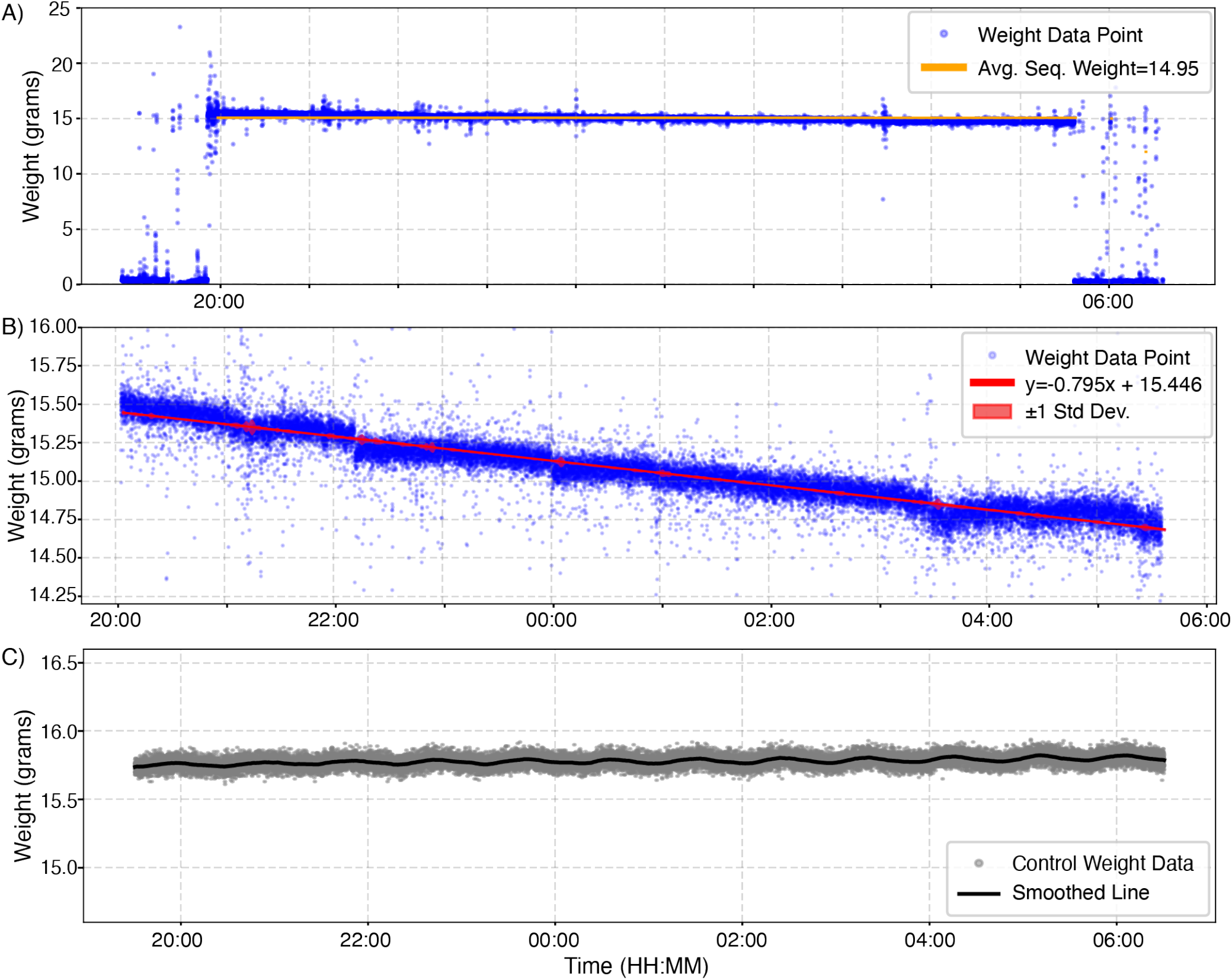
Linear regression of canary weight overnight. **(A)** Weight data measurements (grams, blue dots) of one canary through darkness hours. The vertical orange line shows the average value calculated from the whole ‘on-scale’ sequence (8PM -5:36AM). **(B)** A closer look at the ‘on-scale’ sequence, where the data points (blue dots) are matched with a linear regression line (red line). The opaque red cloud around the red line shows *±*1 standard deviation for every data point (calculated as the standard deviation of a rolling window with a window size of 200). This line depicts the regression of the canary’s weight during the night period (0.08 g, or 0.52% of bodyweight loss per hour, for a total of 0.8 g weight loss, which is 5.17% of its bodyweight). **(C)** Results from the control experiment of weighing an idle object during night time. Measured data points (grams) and smoothed line (grey dots and black line accordingly) show little-to-no fluctuations in the measured weight (*±*0.05 g). This validates that the measured canary weight regression overnight is not a result of scale deviations.

We also tried to extract another estimation of the canary’s weight in a more robust method. First, we took the entire dataset and cleaned it from baseline noise (the signal of all ‘off-scale’ measurements and other fluctuations that could have been measured by occasional touches or even wind generated by the canary’s wing flaps). Here we used 2 g as the threshold. We then binned the data with a bin size of 0.1 g and found the most frequent measurements. Fig. 3 shows one example of this estimation, where the most frequent measurement is 17.88 g, and the mean value is 17.05 g.

Further analysis of the data from two canaries exposed hours-long sequences of consecutive weight measurements during darkness hours, indicating that canaries were indeed sleeping on the scale. From this unique data we were able to measure linear regression of the weight loss in that period. Fig. 4 shows one example for the bird weight over time during over 9 hours of standing on the scale, along with the mean weight (Fig. 4A) and the same data with a linear regression fit line (Fig. 4B). With a start weight of 15.48 g at the beginning of the night and an end weight of 14.68 g in the morning, the bird lost a total of 0.8 grams over a period of nearly 9.5 hours, which is a total of 5.17% of its body weight. Per hour, the bird lost 0.08 g which is 0.52% of its body weight. Such results were recorded 3 times, and all of them suggest the same pattern -demonstrating that our system is sensitive enough to measure weight changes below 1% of the body weight of a 15g canary. To further validate these findings and eliminate the possibility that the decrease in weight is a result of scale deviations, we tested the scale system in similar conditions. A pair of tweezers with known weight (17.3 g) was put on the scale for 12 hours consecutively, from which we extracted the data between the onset and offset times of the canary on the scale overnight. Light conditions inside the acoustic chambers were similar to the ones the canaries experienced. The results show little-to-no fluctuations that could explain the decrease we observed in the real canary overnight data (Fig. 4C). Put together, these results clearly demonstrate natural weight loss overnight in canaries, and a part of normal daily weight fluctuations [8] that has never been demonstrated in neuroscience experimental setups.

## 5 Discussion

Automated weighing systems have been developed for different laboratory species, such as mice, but until now, there has been no equivalent system designed specifically for birds in neuroscience experiments. Monitoring the weight of research birds could benefit their wellbeing as well as ensure reliable scientific data collection, as body weight is a key marker of their health [10]. Traditional weighing methods, like manually placing birds in cloth bags [5], are disruptive and can cause stress, which may interfere with natural behaviors like singing. Our novel automated system addresses these limitations by enabling continuous, hands-free weight monitoring, ensuring accurate health assessments without disrupting the birds’ natural behavior.

Our results demonstrate the effectiveness of this system in capturing valuable, real-time data on both tethered and untethered birds. One notable finding was the consistent pattern of overnight weight loss. In two of the birds, where we were able to calculate a linear regression to track weight loss percentages during periods of extended inactivity. For example, we observed one bird losing 0.8 grams, or approximately 5.17% of its body weight, over a 9.5-hour period of sleep. This ability to continuously monitor weight allowed us to track such physiological changes with high precision, offering insights into daily patterns that might otherwise go unnoticed in typical manual weighing methods. Looking ahead, this system could serve as a foundation for more elaborate studies that monitor birds under different conditions and research paradigms. For instance, future research could use this system to track how environmental factors, changes in diet, or hormonal cycles influence weight fluctuations over time. The non-intrusive nature of our method makes it particularly suitable for long-term studies, where continuous data collection is essential for understanding how birds adapt to varying experimental conditions.

However, while our system advances on existing methods to monitor birds, it is only the first step in establishing a reliable approach to improving the care for birds in longitudinal experiments. Specifically, as was done in the case of using singing rate to assess stress [15], weight measurements and their dynamics need to be linked to changes in health, stress, and well being of birds. Closing this gap will require a significant data collection effort going forward. By providing the complete specifications, design files, and software of our system we hope to help initiate this effort.

Furthermore, weight alone is only one parameter for assessing well-being. Future efforts could also focus on integrating other health metrics, such as body condition scoring (BCS) [14], to provide a more comprehensive picture of animal welfare. As seen in other species, combining multiple non-invasive indicators can enhance our ability to monitor the health of research animals, providing a better understanding of their well-being during experimental studies [1].

As mentioned in the Methods, in our experiment the birds had perching options other than the scale, and from two of them (one third) we were not able to gather substantial weight data. Future studies should consider keeping the scale as the only perch available, and provide access to food and water accordingly. Furthermore, cleaning and maintaining of the cage while the scale is installed needs to be handled with care, and re-calibrating the scales following maintenance might be a necessary procedure.

In conclusion, our automated weighing system offers researchers a reliable, non-invasive tool for continuous weight monitoring. This system not only reduces the stress associated with traditional weighing methods but also paves the way for more comprehensive studies that can further explore the physiological responses of birds to a range of experimental conditions.

## 6 Supplementary Data

All supplementary materials can be found here:

https://github.com/NeuralSyntaxLab/Bird-Scale-Methods-Article

Materials includes: 1. Scale design files 2. Scale setup and assembly guides 3. Full parts list 4. Arduino codes 5. Python code for data acquisition

